# Oxygen Microenvironments in *E. coli* Biofilm Nutrient Transport Channels: Insights from Complementary Sensing Approaches

**DOI:** 10.1101/2024.07.20.603676

**Authors:** Beatrice Bottura, Gail McConnell, Lindsey C. Florek, Marina K. Smiley, Ross Martin, Ash Eana, Hannah T. Dayton, Kelly N. Eckartt, Alexa M. Price-Whelan, Paul A. Hoskisson, Lars E.P. Dietrich, Liam M. Rooney

**Author notes:** Co-first Authors (Beatrice Bottura, Gail McConnell).

## Abstract

Chemical gradients and the emergence of distinct microenvironments in biofilms are vital to the stratification, maturation and overall function of microbial communities. These gradients have been well characterised throughout the biofilm mass but the microenvironment of recently discovered nutrient transporting channels in *Escherichia coli* biofilms remains unexplored. This study employs three different oxygen sensing approaches to provide a robust quantitative overview of the oxygen gradients and microenvironments throughout the biofilm transport channel networks formed by *E. coli* macrocolony biofilms. Oxygen nanosensing combined with confocal laser scanning microscopy established that the oxygen concentration changes along the length of biofilm transport channels. Electrochemical sensing provided precise quantification of the oxygen profile in the transport channels, showing similar anoxic profiles compared with the adjacent cells. Anoxic biosensing corroborated these approaches, providing an overview of the oxygen utilisation throughout the biomass. The discovery that transport channels maintain oxygen gradients contradicts the previous literature that channels are completely open to the environment along the apical surface of the biofilm. We provide a potential mechanism for the sustenance of channel microenvironments via orthogonal visualisations of biofilm thin sections showing thin layers of actively growing cells. This complete overview of the oxygen environment in biofilm transport channels primes future studies aiming to exploit these emergent structures for new bioremediation approaches.

## Introduction

The spatial and structural heterogeneity of biofilms leads to the formation of complex emergent architectural features that maintain their durability and ability to distribute key resources. Recently discovered *E. coli* biofilm transport channels present an example of such structures^1^. These nutrient transport channels were discovered using novel mesoscopic imaging methods^2^. They were found to form according to their local environmental conditions^3^ as an emergent property governed by cell shape and driven by cell-to-cell and cell-surface interactions^4^. The proven ability of these channels to translocate nutrients and small particles within mature macrocolony biofilms presents a timely opportunity to develop exploitative channel-mediated bioremediation strategies to lower the impact of biofilms on public health and industry. However, the chemical microenvironment of biofilm transport channels must be elucidated prior to the development of new chemical treatments, to inform their design and delivery.

Chemical gradients in biofilms are well characterised and play a key role in nutrient cycling, waste product management, secretion of secondary metabolites and signalling compounds, and oxygen, redox, and pH homeostasis^5–9^. The oxygen profile of mature biofilms is perhaps the most important of these when designing new combative biofilm treatments. There are three key reasons for this: the potential for free and reactive oxygen species (ROS) to degrade the molecular structure of antimicrobial agents^10,11^, the synergistic role that free oxygen and ROS may play in potentiating biofilm destruction when combined with antimicrobial treatments^12,13^, and the emergence of antimicrobial tolerance fuelled by oxygen depletion in mature biofilms^14–17^. Therefore, an understanding of the oxygen gradients and microenvironment of biofilm transport channels is important as it will inform the development of future mitigation strategies which may seek to exploit such channels as drug delivery pipelines.

This study aims to elucidate the oxygen gradients and microenvironment of *E. coli* biofilm transport channels using a multimodal approach to oxygen sensing (Figure 1). We present three complementary sensing approaches to probe the oxygen profile of these structures, and we conclude with a potential structural mechanism for maintaining the homeostasis of oxygen gradients within the channel networks. Based on our previous work, we initially hypothesised that the channel structures formed akin to chasms; open to the air, permitting free diffusion of atmospheric oxygen into the biofilm channels. However, this study determined that the oxygen microenvironment of biofilm transport channels was depleted of oxygen, with a similar oxygen profile to the biofilm cell mass and maintained by a thin apical layer of cells at the crest of each channel structure. Bioremediation strategies can now be developed that capitalise on these observations and our robust overview of the oxygen gradients throughout these channel networks.

**Figure 1.**
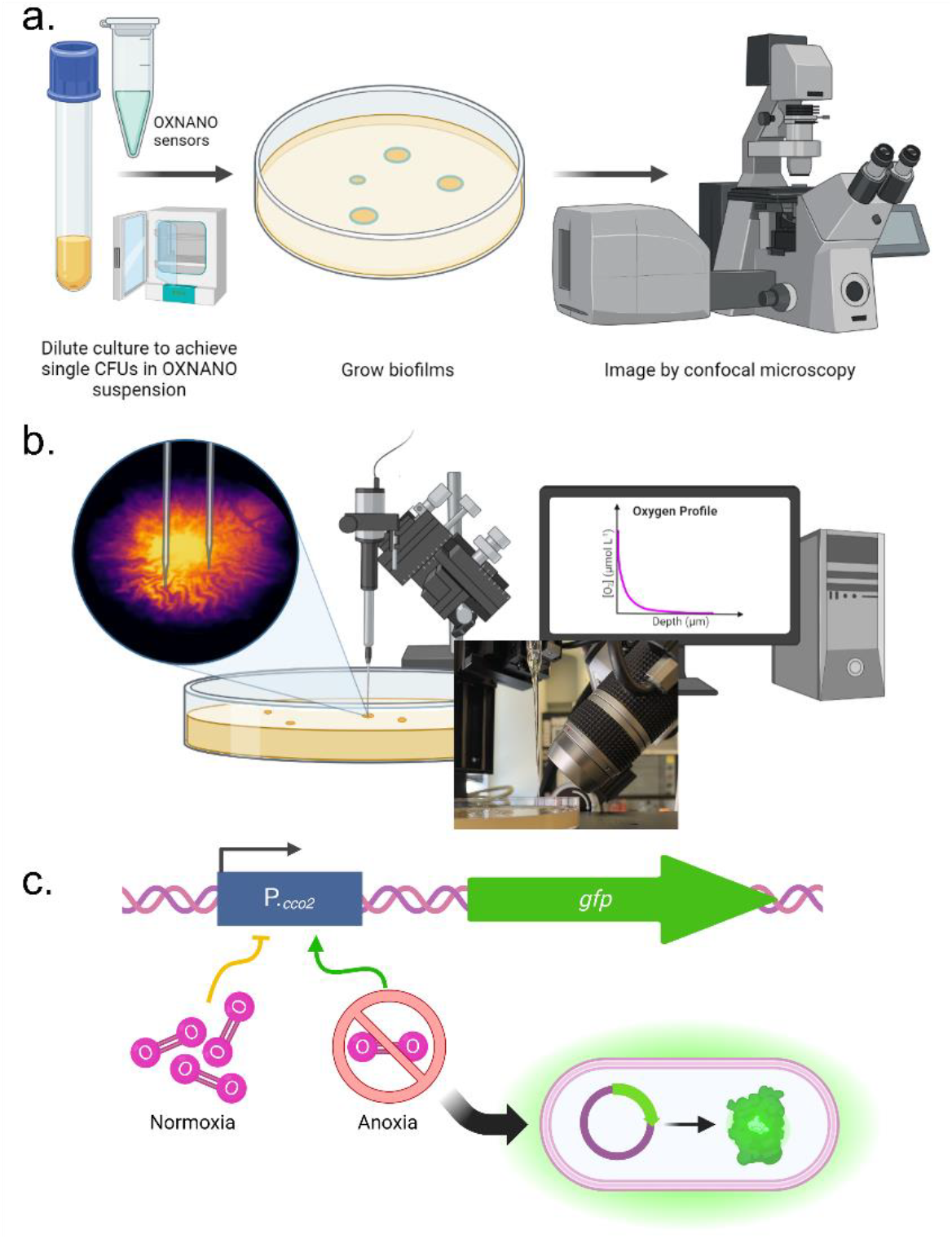
A combinatorial approach to oxygen sensing in established biofilms. **(a)** Oxygen nanosensing uses fluorescent nanospheres which increase in fluorescence intensity proportionally to the local oxygen concentration, providing a ratiometric intensity-based readout of oxygen concentration in complex samples. **(b)** Oxygen sensing via partial pressure measurements achieved using fine-tipped electrochemical probes permits targeted and absolute measurements of the oxygen concentration in different sub-populations of the biofilm by guiding the sensor tip to different regions, with the option of measuring through biofilm depth using a micromanipulator. **(c)** Oxygen biosensing provides insight into the local oxygen environment at the cellular level by directly sensing the cellular response to environmental oxygen via a transcriptional fusion of *gfp* to the promotor of the terminal cytochrome c oxidase, *cco2*.

## Results

### Oxygen Nanosensing Reveals the Oxygen Gradients along Biofilm Transport Channels

Biofilm transport channels were observed to transport oxygen-sensing nanoparticles, thereby providing means to quantify the oxygen microenvironment within channels in comparison to the external surroundings. Figure 2 shows the uptake and modulation of the fluorescence emission intensity of OXNANO sensors throughout a HcRed1-expressing JM105 biofilm. A polar transform facilitated the projection of radially propagating channels as linear structures with a clear boundary at the periphery of the biofilm (Figure 2a). Internalised nanosensors exhibited an 86% decrease in the median relative fluorescence intensity compared to the external population exposed to atmospheric oxygen concentrations, indicating that the oxygen concentration was being attenuated along the length of the transport channels. While these data confirmed the change in the oxygen microenvironment, they do not provide absolute quantification of the oxygen concentrations within these emergent structures.

**Figure 2.**
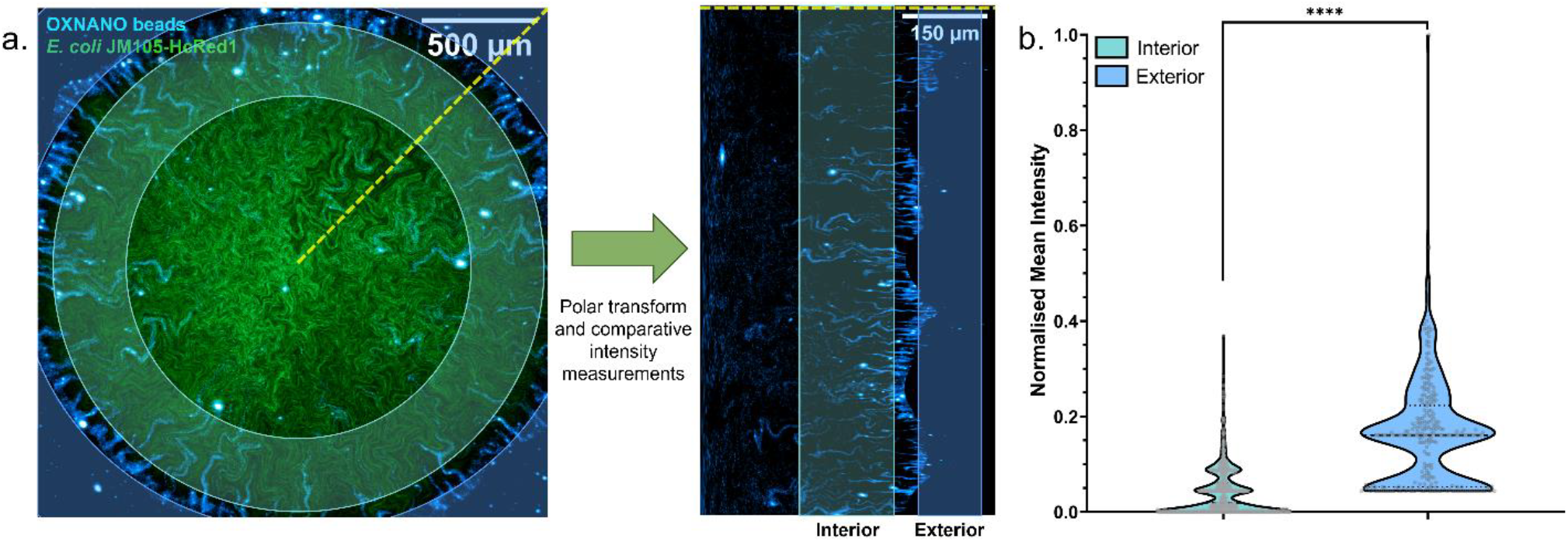
Oxygen Nanosensing Reveals the Oxygen Gradients along Biofilm Transport Channels. **(a)** A maximum intensity projection of an *E. coli* biofilm (green), showing dark transport channel networks, merged with fluorescent oxygen-sensing nanoparticles (cyan). Comparative intensity analysis between the exterior and interior nanosensors was conducted by performing a *polar transform* operation and segmenting nanoparticles from each region for analysis. **(b)** Nanosensors in biofilm channels exhibited an 86% reduction in median fluorescence intensity compared to exterior nanosensors, indicating a decrease in oxygen concentration along the channel network towards the core of the biofilm (p <0.0001, ****) (N_External_ = 270, N_Internal_ = 786; acquired over three biological replicates).

Calibration of the OXNANO beads was attempted by imaging under the same acquisition parameters as biofilm specimens but using lawns of beads either exposed to the atmosphere or mounted and sealed in an anoxic ascorbic acid buffer (Supplementary Figure 1). However, the calibration data did not provide a sufficient difference in the emission intensity between the two calibration points and therefore positioned OXNANO data as purely ratiometric.

### Quantitative Electrochemical Oxygen Sensing Provides a Direct Readout of the Oxygen Gradients throughout Biofilm Transport Channels

Electrochemical oxygen profiling was used to provide a high-resolution readout of the oxygen microenvironment over the depth of biofilm transport channels and compared to the adjacent interstitial cell population. Figure 3a illustrates the careful guidance of a 10 µm-tip oxygen sensing electrode directly into the apical surface of a transport channel and the adjected cell population. The atmospheric oxygen concentration was measured as 266.3 µmol/L at a temperature of 23.3°C during calibration. Figure 3b shows the oxygen concentration through the depth of the biofilm transport channels for wild-type, *gfp*, and *HcRed1*-expressing biofilms. The oxygen concentration decreased immediately under the surface of all biofilms, reaching 0.3 µmol/L within 20 µm of the apical surface. The anoxic region persisted until the basal layers of the biofilm, where partial recovery in the oxygen concentration was observed in all strains. However, the channels consistently exhibited a higher recovery in the oxygen concentration in these regions compared to the interstitial cell populations. The wild-type strain exhibited routinely lower oxygen recovery levels compared to the fluorescent strains.

**Figure 3.**
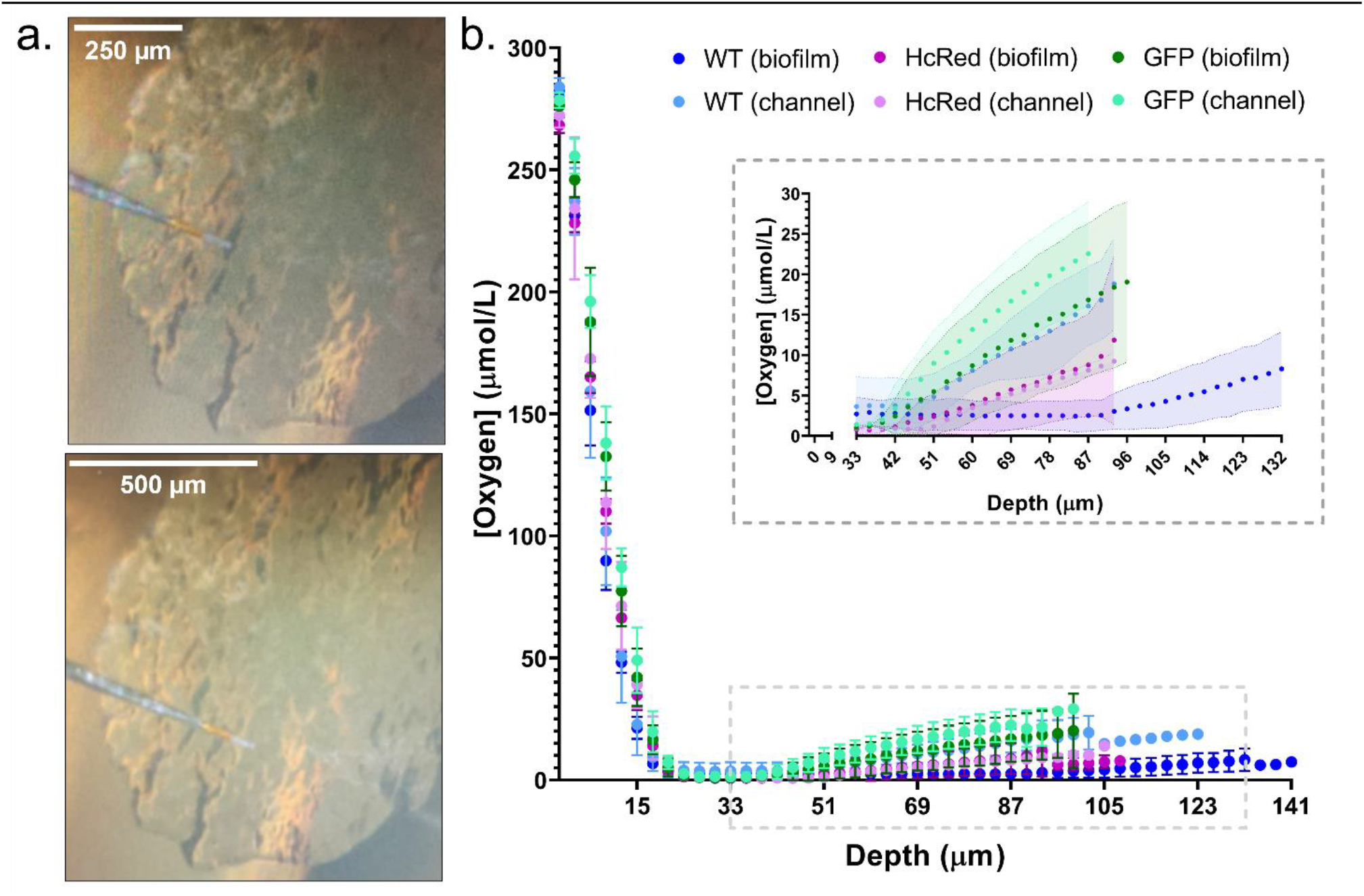
Oxygen concentrations decrease through biofilm depth and differ at the biofilm base. **(a**) Images showing the insertion of an oxygen microelectrode probe into a biofilm transport channel (top) and biofilm cell population (bottom). **(b)** A measurement trace of the oxygen profile through the depth of the biofilm in channel and cell populations showing the mean oxygen concentration and standard deviation for each measurement position. Both channel and biofilm populations exhibited a decrease in oxygen concentration from atmospheric oxygen (266 µmol/L) to 0.3 µmol/L within 20 µm from the biofilm surface. Transport channels exhibited higher oxygen concentrations at depth compared to the adjacent cell populations. A cropped inset is presented showing oxygen recovery at the basal portion of each population. The non-fluorescent wild-type (blue), GFP (green), and HcRed1-expressing (magenta) strains were measured, N = 3 for each condition and population.

### Oxygen Biosensing Concurs with Nanosensing and Electrochemical Sensing to Provide a Robust Overview of Biofilm Transport Channel Oxygen Microenvironments

Figure 4 shows a maximum-intensity projection of a biofilm expressing a fluorescent P_*cco2*_::*gfp* oxygen reporter that conditionally expresses *gfp* under anoxic conditions with high fidelity. These data show an anoxic core accompanied by diffuse fluorescence emission along the transport channel networks with the highest intensity at the centre of the biofilm and a gradual decrease towards the leading edge, interspersed with anoxic subpopulations matching the spatial pattern of canonical biofilm transport channels reported elsewhere^1,3,4,9^. The spatial overview of oxygen gradients from these data concurs with the nanosensing and electrochemical profiling data, which show a change in oxygen gradients along the channel structures and similarities between the microenvironments of the channels and interstitial cell populations, respectively.

**Figure 4.**
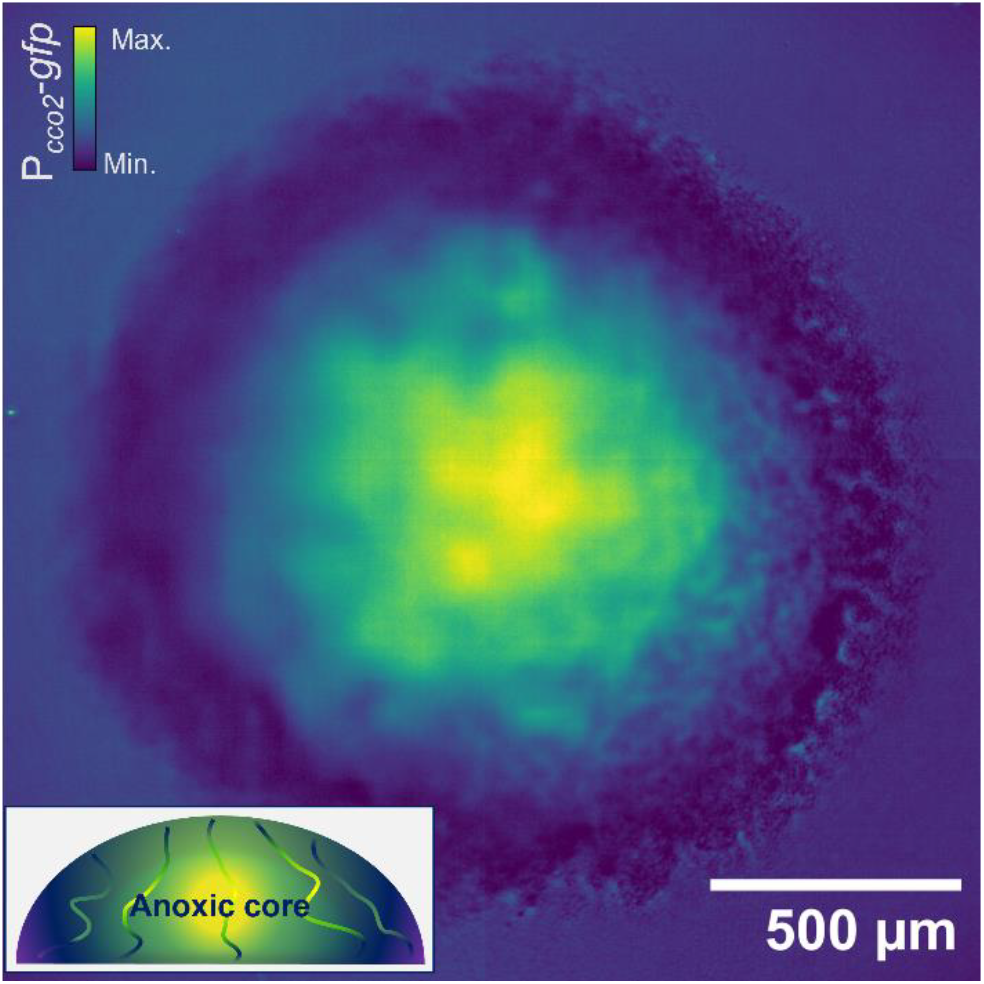
Oxygen biosensing confirms the oxygen profile of biofilm transport channels. A maximum intensity projection of a mature *E. coli* biofilm expressing a *cco2* promoter-*gfp* fusion (P*cco*_*2*_-*gfp*). The emission intensity of GFP indicates levels of anoxia throughout the biofilm, commensurate with a dense hypoxic core. Diffuse signal from interstitial cell populations surrounding the channel structures can be observed permeating radially from the centre. A schematic *x, z* cross-section illustrates the anoxic gradients shown by the oxygen sensing, which concur with nanosensing and electrochemical sensing methods.

### Thin Sectioning Suggests a Mechanism for Maintaining Oxygen and Nutrient Gradients in Biofilm Transport Channels

Our data show that the overall oxygen concentration does not significantly differ between the lumen of the transport channels and the biofilm cell population. This result confounded the original proposed structure of biofilm transport channels, which assumed that they were open to the environment thereby permitting free diffusion of atmospheric oxygen into the lumen of the channels. However, by imaging thin sections of macrocolony biofilms, we have determined that the apical surface of the biofilm remains sealed by a thin layer of capping cells at all points (Figure 5a). Even at positions where transport channels are observed, they are not open to the atmosphere as first presumed from lateral viewpoints. We propose that this layer of actively respiring cells and matrix components is sufficient to maintain the oxygen gradients we observe via nanosensing, electrochemical sensing and biosensing methods. Moreover, we observed that the inferior portions of biofilm transport channels are often open to the underlying substrate. This observation supports the nutrient transport findings documented in other studies^1,3^ and provides an explanation for the increase in oxygen recovery in biofilm channels revealed from our electrochemical sensing data. Figure 5b provides a diagrammatic representation summarising a working model of how biofilm transport channels maintain a marked oxygen gradient, partial recovery from the diffusion of free oxygen and acquisition of nutrients from their underlying grown medium substrate.

**Figure 5.**
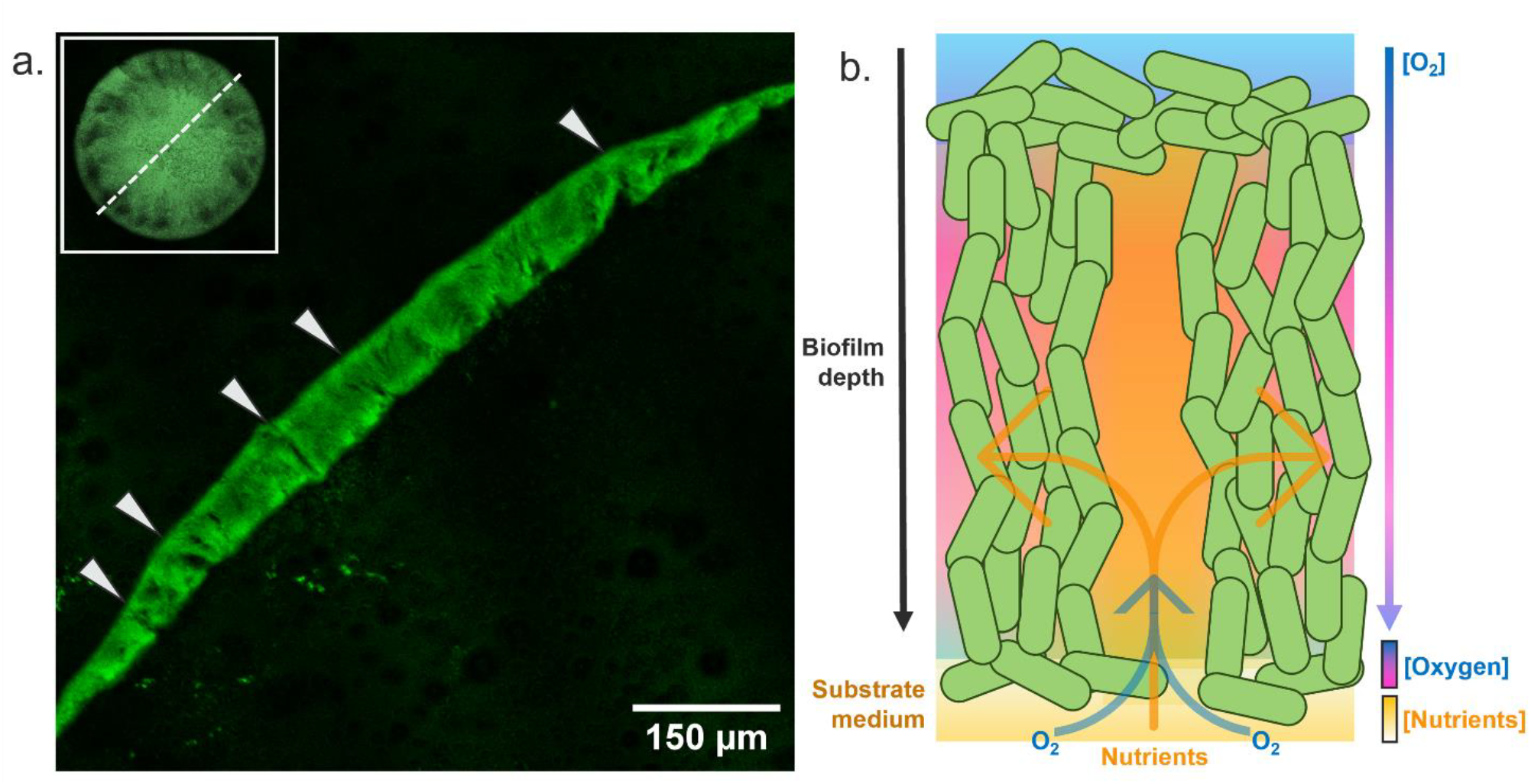
Orthogonal visualisation presents an explanation for chemical gradients in biofilm transport channels. **(a)** A thin section of a fixed *gfp*-expressing *E. coli* JM105 biofilm is presented with arrows at the apical surface indicating regions where transport channels can be viewed sagittally. Thin layers of cells cap the apical surface of the biofilm and enclose the top of transport channels. The inferior portion of the channels remains open to the underlying substrate. A representative view of the section position through the centre of the colony is presented (top left) **(b)** A schematic proposing nutrient and oxygen gradients through the depth of biofilm transport channels. A thin layer of apical cells seals the top of channels and maintains a strict oxygen gradient while nutrients and free oxygen can diffuse from the basolateral surface through open channel conduits into the biomass.

## Discussion

The oxygen profile and chemical microenvironment of biofilm transport channels have remained unexplored since their discovery. This study provides three complementary approaches to quantify the oxygen microenvironment of biofilm channels to provide insight for follow-on studies that might exploit these structures for bioremediation purposes. We used a combination of oxygen nanosensing, electrochemical sensing and biosensing to provide a robust measurement of oxygen gradients throughout biofilm channel networks. These data show that, while oxygen gradients change along the length of biofilm transport channels, there is a sharp oxygen gradient along their depth. Images of biofilm thin sections confirm that biofilm transport channels are capped by a thin layer of cells covering the apical surface of the channel structures while they remain open to the underlying growth substrate. This observation provides a potential mechanism for maintaining such chemical gradients within biofilm transport channels.

Key resource trafficking by biofilm transport channels has been well documented in recent literature^1,3,4,18^, but it remained unclear if these seemingly devoid structures were either hypaethral or enclosed and, thereby, the chemical microenvironment was unknown. The channel structures we discuss here were originally hypothesised to be open chasms spreading along the radii of macrocolony biofilms due to their appearance as dark voids during optical imaging experiments. However, this study concluded that thin layers of surface cells seal and maintain the chemical gradients within the biofilm channels, unlike other larger scale water channels reported in mushroom-shaped *P. aeruginosa*^19,20^, *Klebsiella*^21^ and *Bacillus* spp.^22,23^ biofilms.

Previous studies have used nanosensors^24^, electrochemical profiling^25–27^ and P_*cco2*_-mediated biosensing^25,28^ to study the oxygen gradients and redox profile of *P. aeruginosa* biofilms. While these studies have provided robust insights into the chemical microenvironment within the bulk mass of pseudomonad biofilms, the void structures of *E. coli* transport channels presented a priority for this study which would inform if the local conditions might impact on future channel-exploiting treatments.

Previous optical oxygen nanosensing techniques have been achieved in planktonic cultures^29^, small *Pseudomonas* spp. aggregates^30,31^, and antimicrobial susceptibility screens^32^. Flamholz *et al*. recently documented the common pitfalls in oxygen nanosensing applications in biofilms, where the constituent cells can produce quenching molecules that may impact on the reliability of optical oxygen quantifications^24^. We therefore provided a spatial overview of the relative change in nanosensor intensity as a function of oxygen depletion along biofilm transport channels and supplemented this ratiometric data with high resolution electrochemical sensing experiments. Other studies have also used optical means to study the chemical microenvironments of microbial communities, such as by transient state monitoring coupled with single plane illumination microscopy^33^, ratiometric oxygen-responsive dye measurements^34^, or planar optodes or *in situ* imaging probes^35^. Each of these techniques are limited either by their spatial resolution, attainable field-of-view and low-throughput, or inhomogeneous illumination, leading to spurious quantification^36^. Oxygen nanosensing coupled to conventional low-power confocal microscopy provides some means of circumventing these challenges, although the fluorescent nanosensor approach requires calibration and supplementary methods to provide reliable quantitative measurements^24^. Overall, this approach requires complementation with secondary methods to verify the nanosensing outputs.

Electrochemical sensing of oxygen gradients using Clark-type sensors has been well documented in microbial communities with many examples^37^ spanning environmental microbiology^38,39^, model systems^40^, pathogen biology^15,41^, and wastewater management^42^. The accuracy of this sensing method, both in terms of precision targeting of biofilm sub-populations using the micron-scale sensor tip and the sensitivity and dynamic range, provide a robust quantitative method for complex microbial communities^43^. The complete oxygen profiles of *E. coli* biofilm transport channels we generated concluded that, although the oxygen environment does change through the depth of the transport channels, it does not differ greatly from the profile of the surrounding biofilm mass. This is important to understand due to the leading role that free molecular oxygen and ROS can play in the destruction of antimicrobial agents^10,11^, in turn, leading to diminished treatment efficacy. Moreover, understanding the chemical microenvironment is important for developing ROS-mediated antimicrobial combination therapies^12,44,45^ and understanding the impact of anoxia-induced antimicrobial tolerance^46–48^. It is unclear why the concentration of oxygen recovers to a greater extent in photoprotein-expressing biofilms compared to the wild-type, but this may be linked to the requirement for free oxygen during chromophore maturation for both GFP^49,50^ and HcRed^51^, or due to the increased thickness observed of wild-type biofilms compared to their fluorescent counterparts.

Biosensing using the P_*cco2*_ reporter system facilitated a direct readout of the oxygen environment in different sub-populations of the biofilm. In this system, the *cco2* promoter drives the expression of *gfp* in response to anoxic conditions at the cellular level, providing higher spatial sensitivity than electrochemical sensing can afford and a simultaneous overview of the entire biofilm using the Mesolens. Previous studies have shown the high fidelity of this biosensor in *P. aeruginosa* biofilms^25,28^, and demonstrated that homologous transcription factors drive the anoxic response in *P. aeruginosa* and *E. coli*^52^, priming the translation of the biosensor for visualising oxygen gradients in *E. coli* biofilms. Our data revealed the canonical anoxic core of the macrocolony biofilms that other studies have reported^5,7,8,53–56^, but also show diffuse signal emitted from channel-lining cells throughout the biofilm, thereby complementing our nanosensing and electrochemical sensing experiments. Finally, the orthogonal visualisation of transport channels using thinly sectioned biofilm specimens revealed a thin crowning layer of cells, where previously the channels were hypothesised to be open to the atmosphere. Previous studies have determined that the generation of biofilm oxygen gradients is not caused by physical barriers (i.e., cells, matrix, etc.), but instead penetration is limited by the active respiration of apical cell layers^56–58^. We therefore propose that the thin layer of cells that we observe is actively contributing to the underlying chemical microenvironment of the channel structures, maintaining the anoxic environment that adjacent fermentative cell populations^6,59–61^ are also exposed to.

The combined outcome from the multimodal oxygen sensing presented in this study provides a robust overview of the oxygen gradients and microenvironments of *E. coli* biofilm transport channels. In turn, this work primes future studies exploring the basic physiology and translational potential of biofilm transport channels for new mitigation strategies.

## Methods

### Strains and Growth Conditions

All experiments were performed using the *E. coli* strains outlined in Supplementary Table 1. Macrocolony biofilms were grown by spreading 100 µl of a 1×10^4^ cfu/ml inoculum on solid LB (Miller) medium, supplemented where appropriate with the required selective antibiotic. Biofilms were grown at 37°C in darkened conditions for 18-24 hours before imaging. Planktonic cultures were maintained using LB (Miller) broth and incubated at 37°C while shaking continuously at 225 rpm.

### Specimen Preparation for Imaging

Macrocolony biofilms were prepared for imaging by inoculating lawns of dilute liquid culture, as above, onto moulds filled with solid growth media. 3D printed chamber moulds were used for imaging applications, as described elsewhere^1,3^. To ensure exposure to atmospheric conditions, biofilms were imaged under air immersion throughout this study; imaging of oxygen nanosensor-supplemented biofilms was performed in air immersion using an inverted confocal laser scanning microscope, as described later, and oxygen biosensor imaging was performed using the biofilm specimen preparation methods described by Baxter *et al*.^18^. All experiments were conducted in triplicate.

### Oxygen Nanosensing in Biofilms

Oxygen nanosensing was conducted using proprietary oxygen-sensing nanoparticles, OXNANO (PyroScience GmbH, Germany). Biofilms were prepared for nanoparticle uptake as previously described^1^. Briefly, a suspension of mid-log phase *E. coli* JM105 miniTn7::*HcRed1* at a density of 1×10^4^ cfu/ml was prepared and supplemented with a final concentration of 10 µg/ml OXNANO particles. A 100 µl aliquot of the nanoparticle culture suspension was inoculated to form discrete microcolony biofilms as described above.

Following growth, specimens were imaged using a Leica SP5 confocal laser scanning microscope (Leica, Germany). Fluorescence excitation for HcRed1 and OXNANO particles was provided via the 543 nm and 633 nm lines of a helium-neon laser, respectively. Fluorescence emission was detected simultaneously using two photomultiplier tubes (PMT) with spectral detection set from 650-690 nm for HcRed1 and 740-780nm for OXNANO particles. Images were acquired using a 10x/0.4NA objective lens (Leica, Germany).

Nanoparticle calibration datasets were acquired using OXNANO bead preparations on LB agar pads imaged under atmospheric exposure or sealed in an anoxic ascorbic acid buffer. OXNANO bead preparations were created by supplementing a 10 µg/ml suspension of beads over a thin LB pad constructed using a 10×10 mm Gene Frame (ThermoFisher Scientific, USA), as described elsewhere^62^. The beads were left to dry onto the pads and imaged as described above while exposed to the air (i.e., without a coverglass), or mounted in an anoxic ascorbic acid buffer solution (10 mM ascorbic acid, 90 mM NaOH) and sealed with a coverglass (Supplementary Figure 1).

### Oxygen Profiling in Biofilms

A 10 µm-tip oxygen microsensor (OX-10; Unisense, Denmark) was used to measure the oxygen concentration throughout the depth of biofilm transport channels and the adjacent interstitial cell population. The microsensor electrode was calibrated using a two-point calibration method. Atmospheric calibrations were conducted using a calibration chamber (CAL300; Unisense, Denmark) containing water bubbled continuously with air for at least 30 minutes before reading. Anoxic calibrations were conducted by submersion of the microsensor tip into an anoxic ascorbic acid buffer solution (10 mM ascorbic acid, 90 mM NaOH) and incubated at room temperature for 30 minutes to acclimatise before registering the ‘zero’ value. The electrode was then cleaned with and stored in distilled water until required.

Macrocolony biofilms grown for 20 hours on LB agar plates were placed on top of an aluminium foil surface so that ambient light was reflected through the base of the biofilm channels and rendered them visible using a VHX-1000 digital microscope (Keyence, Japan) set on a rotational mount to 45°. The microsensor electrode was mounted on a micromanipulator (MM33; Unisense, Denmark) to facilitate automated stepped movement of the electrode through the depth of the biofilm. Profiles were recorded using a multimeter and the SensorTrace Profiling software (Unisense, Denmark), as described by Jo *et al*.^25^. This setup provided means to measure the oxygen concentration through the depth of a given transport channel by incrementally stepping the electrode from the apical to the basolateral surface of the biofilm. Readings were acquired with a measurement time of 3 seconds and a dwell time between measurements of 5 seconds. A step size of 5 µm was used to measure the entire depth of the biofilm (typically ranging from 100 µm – 150 µm). Both the channel and interstitial regions were measured in triplicate for each strain.

### Oxygen Biosensing in Biofilms

The oxygen reporter plasmid, pAW9 (Supplementary Table 1)^28,52^, was transformed into JM105 via electroporation and maintained during all subsequent experiments using 100 µg/ml ampicillin. Macrocolony biofilms were grown as described for imaging above and imaged using the Mesolens in widefield epifluorescence mode, as described previously^1,3^. Briefly, a 490 nm LED (pE-4000; CoolLED, UK) provided an excitation source for GFP expressed under anoxic conditions. Fluorescence emission was captured using the Mesolens coupled to a VNP-29MC CCD camera with a chip-shifting modality (Vieworks, South Korea) with a triple-bandpass filter (540 ± 10 nm) placed in the detection pathway. Mesoscopic imaging was conducted using water immersion (*n* = 1.33) with the correction collars set accordingly to minimise spherical aberration via refractive index mismatch.

### Paraffin-embedding and Biofilm Thin Sectioning

Thin sections of macrocolony were generated and imaged by widefield epifluorescence microscopy to provide a sagittal view of biofilm transport channels (Supplementary Figure 2). Biofilms were prepared according to the methods described by Cornell *et al*.^63^ and Smiley *et al*.^27^. Briefly, JM105 miniTn7::*HcRed1* biofilms were grown as described above and prepared for processing by embedding in cooled molten 1% (w/v) agarose. 15 ml of 1% agarose was delicately poured over the surface of the Petri dish containing macrocolony biofilms. After setting, each was excised with a clear margin surrounding the biofilm as a cube of solid agar base and agarose top. Specimens were placed into histocassettes and fixed overnight in 4% (w/v) paraformaldehyde in phosphate-buffered saline (PBS) (pH 7.2) before washing in increasing ethanol concentrations (25%, 50%, 70%, 95% in PBS and three times in 100% ethanol). The biofilms were then cleared with three washes of Histo-Clear II (National Diagnostics, USA), infiltrated with molten paraffin wax (Leica Biosystems, Germany), and left to solidify for three hours. Biofilms were then mounted and sectioned into 10 µm slices using an automatic microtome (905200ER; ThermoFisher Scientific, USA). Specimens were mounted on clean glass slides and air died overnight before heat fixing at 45°C for 30 minutes. The slides were rehydrated in PBS by decreasing through the ethanol gradient above and mounted in ProLong™ Diamond Antifade Mountant (Invitrogen, USA) before being sealed with Type 1.5 coverglasses. Specimens were imaged using an inverted IX81 microscope coupled to a FluoView FV1000 confocal laser scanning microscope (Olympus, Japan). Fluorescence excitation for GFP was provided a 488 nm argon laser (GLG3135; Showa Optronics, Japan). Fluorescence emission was detected using a PMT with spectral detection set from 510-550 nm. Images were acquired using a 10x/0.4NA objective lens (Olympus, Japan).

### Image Analysis

All image processing and analyses were conducted using FIJI v1.54f^64^. Images were false-coloured and linearly contrast-adjusted post-analysis for display purposes where appropriate.

For oxygen nanosensing, images were pre-processed by performing a *polar transform* function^65^ to transform from polar to Cartesian coordinates and simplify the analysis of radially projecting channel structures. The coordinates of the centre of the biofilm were calculated using the *Measure* function in FIJI and the number of pixels per angular coordinates was set to 4096 pixels (11 pixels per degree). Converting to a Cartesian projection of the image facilitated the selective measurement of exterior and internalised nanosensor populations. A region of interest was cropped from the exterior of the biofilm and the internal region, and a median filter (σ = 5 pixels) was used to reduce noise contributions from the analysis with minimal impact on the signal intensity. The nanosensing beads were thresholded using automatic *yen* parameters, and the fluorescence emission intensity of the OXNANO sensors was measured using the *Analyse Particles* function to generate the mean fluorescence intensity of each nanosensor particle.

### Statistical Analysis

Statistical analyses were conducted using Prism v8.0.2. (GraphPad, USA). Following a normality test, the intensity of OXNANO particles between the internal and exterior populations was compared using a two-tailed Mann-Whitney test. The emission intensity data were normalised ranging between the maximum and minimum intensity of the thresholded beads 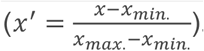 The OXNANO calibration data were also compared using a two-tailed Mann-Whitney test.

## Additional Information

## Acknowledgements

The Authors wish to thank Prof. Marvin Whiteley (Georgia Institute of Technology, GA, USA) for the kind gift of the pAW9 plasmid. BB and RM were funded by the University of Strathclyde. GM was funded by The Medical Research Council, MR/K015583/1, and the Biotechnology and Biological Sciences Research Council, BB/P02565X/1 and BB/T011602/1. AE was funded by a Microbiology Society Harry Smith Vacation Studentship and a Royal Microscopical Society Summer Studentship. PAH was funded by the Royal Academy of Engineering Research Chair Scheme for long-term personal research support (RCSRF2021\11\15). LEPD was funded by the National Institutes of Health and the National Institute of Allergy and Infectious Diseases (R01AI103369). LMR was funded by the Leverhulme Trust, the University of Strathclyde and a Scottish Universities Life Science Alliance Early Career Development Award, 2023.

Figure 1, Supplementary Figure 1 and Supplementary Figure 2 were prepared using BioRender.com (Licence Number: VF272F7XHX).

## Authors’ Contributions

BB: analysis, investigation, methodology, visualisation, manuscript preparation and review.

GM: funding acquisition, analysis, methodology, supervision, manuscript preparation and review.

LCF: analysis, investigation, methodology, manuscript preparation and review.

MKS: investigation, methodology, manuscript preparation and review.

RM: investigation, validation, manuscript preparation and review.

AE: investigation, manuscript preparation and review.

HTD: investigation, validation, manuscript preparation and review.

KNE: investigation, validation, manuscript preparation and review.

AMP: manuscript preparation and review.

PAH: manuscript preparation and review.

LEPD: conceptualisation, funding acquisition, project administration, analysis, methodology, supervision, manuscript preparation and review.

LMR: conceptualisation, funding acquisition, project administration, data curation, analysis, investigation, methodology, supervision, validation, visualisation, manuscript preparation and review.

## Data Availability Statement

The data that support the findings of this study are openly available via the University of Strathclyde KnowledgeBase data repository (doi.org/10.15129/1e4a5f5e-24a3-42fc-997a-585877ccb4d8).

## Competing Interests

We declare no competing interests

## Notes

### Competing Interest Statement

The authors have declared no competing interest.

## References

1. Rooney, L. M., Amos, W. B., Hoskisson, P. A. & McConnell, G. Intra-colony channels in E. coli function as a nutrient uptake system. The ISME Journal (2020) doi:10.1038/s41396-020-0700-9.

2. Rooney, L. M. et al. Addressing multiscale microbial challenges using the Mesolens. Journal of Microscopy jmi.13172 (2023) doi:10.1111/jmi.13172.

3. Bottura, B., Rooney, L. M., Hoskisson, P. A. & McConnell, G. Intra-colony channel morphology in Escherichia coli biofilms is governed by nutrient availability and substrate stiffness. Biofilm 4, 100084 (2022).

4. Bottura, B., Rooney, L., Feeney, M., Hoskisson, P. A. & McConnell, G. Fractal complexity of Escherichia coli nutrient transport channels is influenced by cell shape and growth environment. Preprint at 10.1101/2023.11.29.569150 (2023).

5. Stewart, P. S. & Franklin, M. J. Physiological heterogeneity in biofilms. Nat Rev Microbiol 6, 199–210 (2008).

6. Jo, J., Price-Whelan, A. & Dietrich, L. E. P. Gradients and consequences of heterogeneity in biofilms. Nat Rev Microbiol 20, 593–607 (2022).

7. Rani, S. A. et al. Spatial Patterns of DNA Replication, Protein Synthesis, and Oxygen Concentration within Bacterial Biofilms Reveal Diverse Physiological States. J Bacteriol 189, 4223–4233 (2007).

8. Díaz-Pascual, F. et al. Spatial alanine metabolism determines local growth dynamics of Escherichia coli colonies. eLife 10, e70794 (2021).

9. Dayton, H. et al. Cellular arrangement impacts metabolic activity and antibiotic tolerance in Pseudomonas aeruginosa biofilms. PLOS Biology 22, e3002205 (2024).

10. Pan, M. & Chu, L. M. Adsorption and degradation of five selected antibiotics in agricultural soil. Science of The Total Environment 545–546, 48–56 (2016).

11. Agarwal, P., Singh, N. & Farooqui, A. Impact of antibiotics on agricultural microbiome: emergence of antibiotic resistance bacteria. in Degradation of Antibiotics and Antibiotic-Resistant Bacteria from Various Sources 231–246 (Elsevier, 2023). doi:10.1016/B978-0-323-99866-6.00012-X.

12. Vatansever, F. et al. Antimicrobial strategies centered around reactive oxygen species – bactericidal antibiotics, photodynamic therapy, and beyond. FEMS Microbiol Rev 37, 955–989 (2013).

13. Van Acker, H. & Coenye, T. The Role of Reactive Oxygen Species in Antibiotic-Mediated Killing of Bacteria. Trends in Microbiology 25, 456–466 (2017).

14. Borriello, G. et al. Oxygen Limitation Contributes to Antibiotic Tolerance of Pseudomonas aeruginosa in Biofilms. Antimicrob Agents Chemother 48, 2659–2664 (2004).

15. Kowalski, C. H., Morelli, K. A., Schultz, D., Nadell, C. D. & Cramer, R. A. Fungal biofilm architecture produces hypoxic microenvironments that drive antifungal resistance. Proc. Natl. Acad. Sci. U.S.A. 117, 22473–22483 (2020).

16. Beebout, C. J., Sominsky, L. A., Eberly, A. R., Van Horn, G. T. & Hadjifrangiskou, M. Cytochrome bd promotes Escherichia coli biofilm antibiotic tolerance by regulating accumulation of noxious chemicals. npj Biofilms Microbiomes 7, 35 (2021).

17. Ciofu, O., Moser, C., Jensen, P.Ø. & Høiby, N. Tolerance and resistance of microbial biofilms. Nat Rev Microbiol 20, 621–635 (2022).

18. Baxter, K. J., Sargison, F. A., Fitzgerald, J. R., McConnell, G. & Hoskisson, P. A. Time-lapse mesoscopy of Candida albicans and Staphylococcus aureus dual-species biofilms reveals a structural role for the hyphae of C. albicans in biofilm formation. Microbiology 170, (2024).

19. Drury, W. J., Characklis, W. G. & Stewart, P. S. Interactions of 1 μm latex particles with Pseudomonas aeruginosa biofilms. Water Research 27, 1119–1126 (1993).

20. Kempes, C. P., Okegbe, C., Mears-Clarke, Z., Follows, M. J. & Dietrich, L. E. P. Morphological optimization for access to dual oxidants in biofilms. Proceedings of the National Academy of Sciences 111, 208–213 (2014).

21. Stoodley, P., Lewandowski, Z., & others. Liquid flow in biofilm systems. Applied and environmental microbiology 60, 2711–2716 (1994).

22. Asally, M. et al. Localized cell death focuses mechanical forces during 3D patterning in a biofilm. Proceedings of the National Academy of Sciences 109, 18891–18896 (2012).

23. Wilking, J. N. et al. Liquid transport facilitated by channels in Bacillus subtilis biofilms. Proceedings of the National Academy of Sciences 110, 848–852 (2013).

24. Flamholz, A. I., Saccomano, S., Cash, K. & Newman, D. K. Optical O2 Sensors Also Respond to Redox Active Molecules Commonly Secreted by Bacteria. mBio 13, e02076–22 (2022).

25. Jo, J., Cortez, K. L., Cornell, W. C., Price-Whelan, A. & Dietrich, L. E. An orphan cbb3-type cytochrome oxidase subunit supports Pseudomonas aeruginosa biofilm growth and virulence. 30 (2017).

26. Schiessl, K. T. et al. Phenazine production promotes antibiotic tolerance and metabolic heterogeneity in Pseudomonas aeruginosa biofilms. Nature Communications 10, (2019).

27. Smiley, M. K. et al. MpaR-driven expression of an orphan terminal oxidase subunit supports Pseudomonas aeruginosa biofilm respiration and development during cyanogenesis. mBio 15, e02926–23 (2024).

28. Wessel, A. K. et al. Oxygen Limitation within a Bacterial Aggregate. mBio 5, e00992–14 (2014).

29. Ladner, T., Flitsch, D., Schlepütz, T. & Büchs, J. Online monitoring of dissolved oxygen tension in microtiter plates based on infrared fluorescent oxygen-sensitive nanoparticles. Microb Cell Fact 14, 161 (2015).

30. Jewell, M. P., Galyean, A. A., Kirk Harris, J., Zemanick, E. T. & Cash, K. J. Luminescent Nanosensors for Ratiometric Monitoring of Three-Dimensional Oxygen Gradients in Laboratory and Clinical Pseudomonas aeruginosa Biofilms. Appl Environ Microbiol 85, e01116–19 (2019).

31. Smriga, S., Ciccarese, D. & Babbin, A. R. Denitrifying bacteria respond to and shape microscale gradients within particulate matrices. Commun Biol 4, 570 (2021).

32. Jusková, P. et al. Real-Time Respiration Changes as a Viability Indicator for Rapid Antibiotic Susceptibility Testing in a Microfluidic Chamber Array. ACS Sens. 6, 2202–2210 (2021).

33. Karampatzakis, A. et al. Measurement of oxygen concentrations in bacterial biofilms using transient state monitoring by single plane illumination microscopy. Biomedical Physics & Engineering Express 3, 035020 (2017).

34. Larsen, M., Borisov, S. M., Grunwald, B., Klimant, I. & Glud, R. N. A simple and inexpensive high resolution color ratiometric planar optode imaging approach: application to oxygen and pH sensing. Limnology & Ocean Methods 9, 348–360 (2011).

35. Kühl, M., Rickelt, L. F. & Thar, R. Combined Imaging of Bacteria and Oxygen in Biofilms. Appl Environ Microbiol 73, 6289–6295 (2007).

36. Sankaran, J., Karampatzakis, A., Rice, S. A. & Wohland, T. Quantitative imaging and spectroscopic technologies for microbiology. FEMS Microbiology Letters 365, (2018).

37. Saccomano, S. C., Jewell, M. P. & Cash, K. J. A review of chemosensors and biosensors for monitoring biofilm dynamics. Sensors and Actuators Reports 3, 100043 (2021).

38. Sørensen, K., Řeháková, K., Zapomělová, E. & Oren, A. Distribution of benthic phototrophs, sulfate reducers, and methanogens in two adjacent saltern evaporation ponds in Eilat, Israel. Aquat. Microb. Ecol. 56, 275–284 (2009).

39. Hicks, N. et al. Oxygen dynamics in shelf seas sediments incorporating seasonal variability. Biogeochemistry 135, 35–47 (2017).

40. Guimerà, X. et al. A Minimally Invasive Microsensor Specially Designed for Simultaneous Dissolved Oxygen and pH Biofilm Profiling. Sensors 19, 4747 (2019).

41. Kiamco, M. M., Atci, E., Mohamed, A., Call, D. R. & Beyenal, H. Hyperosmotic Agents and Antibiotics Affect Dissolved Oxygen and pH Concentration Gradients in Staphylococcus aureus Biofilms. Appl Environ Microbiol 83, e02783–16 (2017).

42. Qi, C. et al. Organic carbon determines nitrous oxide consumption activity of clade I and II nosZ bacteria: Genomic and biokinetic insights. Water Research 209, 117910 (2022).

43. Revsbech, N. P. Simple sensors that work in diverse natural environments: The micro-Clark sensor and biosensor family. Sensors and Actuators B: Chemical 329, 129168 (2021).

44. Akbari, M. Z., Xu, Y., Lu, Z. & Peng, L. Review of antibiotics treatment by advance oxidation processes. Environmental Advances 5, 100111 (2021).

45. Qi, W., Jonker, M. J., de Leeuw, W., Brul, S. & ter Kuile, B. H. Reactive oxygen species accelerate de novo acquisition of antibiotic resistance in E. coli. iScience 26, 108373 (2023).

46. Schaible, B., Taylor, C. T. & Schaffer, K. Hypoxia Increases Antibiotic Resistance in Pseudomonas aeruginosa through Altering the Composition of Multidrug Efflux Pumps. Antimicrob Agents Chemother 56, 2114–2118 (2012).

47. Jensen, P.Ø., Kolpen, M., Kragh, K. N. & Kühl, M. Microenvironmental characteristics and physiology of biofilms in chronic infections of CF patients are strongly affected by the host immune response. APMIS 125, 276–288 (2017).

48. Olsen, I. Biofilm-specific antibiotic tolerance and resistance. Eur J Clin Microbiol Infect Dis 34, 877–886 (2015).

49. Reid, B. G. & Flynn, G. C. Chromophore Formation in Green Fluorescent Protein. Biochemistry 36, 6786–6791 (1997).

50. Barondeau, D. P., Putnam, C. D., Kassmann, C. J., Tainer, J. A. & Getzoff, E. D. Mechanism and energetics of green fluorescent protein chromophore synthesis revealed by trapped intermediate structures. Proc. Natl. Acad. Sci. U.S.A. 100, 12111–12116 (2003).

51. Wilmann, P. G. et al. The 2.1Å Crystal Structure of the Far-red Fluorescent Protein HcRed: Inherent Conformational Flexibility of the Chromophore. Journal of Molecular Biology 349, 223–237 (2005).

52. Winteler, H. V. & Haas, D. The homologous regulators ANR of Pseudornonas aeruginosa and FNR of Escherichia coli have overlapping but distinct specificities for anaerobically inducible promoters. Microbiology 142, 685–693 (1996).

53. de Beer, D., Stoodley, P., Roe, F. & Lewandowski, Z. Effects of biofilm structures on oxygen distribution and mass transport. Biotech & Bioengineering 43, 1131–1138 (1994).

54. Depetris, A. et al. Biophysical properties at patch scale shape the metabolism of biofilm landscapes. npj Biofilms Microbiomes 8, 5 (2022).

55. Zhang, Y. et al. A Microfluidic Approach for Quantitative Study of Spatial Heterogeneity in Bacterial Biofilms. Small Science 2, 2200047 (2022).

56. Tokunou, Y., Toyofuku, M. & Nomura, N. Physiological Benefits of Oxygen-Terminating Extracellular Electron Transfer. mBio 13, e01957–22 (2022).

57. Barraud, N., Kelso, M., Rice, S. & Kjelleberg, S. Nitric Oxide: A Key Mediator of Biofilm Dispersal with Applications in Infectious Diseases. CPD 21, 31–42 (2014).

58. Stewart, P. S. et al. Reaction–diffusion theory explains hypoxia and heterogeneous growth within microbial biofilms associated with chronic infections. npj Biofilms Microbiomes 2, 16012 (2016).

59. Serra, D. O., Klauck, G. & Hengge, R. Vertical stratification of matrix production is essential for physical integrity and architecture of macrocolony biofilms of Escherichia coli. Environmental Microbiology 17, 5073–5088 (2015).

60. Létoffé, S. et al. Biofilm microenvironment induces a widespread adaptive amino-acid fermentation pathway conferring strong fitness advantage in Escherichia coli. PLoS Genet 13, e1006800 (2017).

61. Martín-Rodríguez, A. J. Respiration-induced biofilm formation as a driver for bacterial niche colonization. Trends in Microbiology 31, 120–134 (2023).

62. de Jong, I. G., Beilharz, K., Kuipers, O. P. & Veening, J.-W. Live Cell Imaging of Bacillus subtilis and Streptococcus pneumoniae using Automated Time-lapse Microscopy. JoVE 3145 (2011) doi:10.3791/3145.

63. Cornell, W. C. et al. Paraffin Embedding and Thin Sectioning of Microbial Colony Biofilms for Microscopic Analysis. JoVE 57196 (2018) doi:10.3791/57196.

64. Schindelin, J. et al. Fiji: an open-source platform for biological-image analysis. Nature Methods 9, 676– 682 (2012).

65. Donnelly, E. & Mothe, F. Polar Transformer. imagej.net/ij/plugins/polar-transformer.html (2007).

66. Lambertsen, L., Sternberg, C. & Molin, S. Mini-Tn7 transposons for site-specific tagging of bacteria with fluorescent proteins. Environmental Microbiology 6, 726–732 (2004).

