## Supplementary Information for "Oxygen Microenvironments in *E. coli* Biofilm Nutrient Transport Channels: Insights from Complementary Sensing Approaches"

^†^Co-first Author

**Supplementary Material**

**Supplementary Figure 1. Oxygen Nanosensor Range Validation. (a)** An experimental schematic to determine the difference in oxygen nanosensor emission intensity under atmospheric and anoxic conditions. Ascorbic acid buffer acts as a strong reducing agent and sequesters molecular oxygen rapidly once the agar pad is sealed with a coverglass. Atmospheric oxygen conditions were achieved by imaging the exposed pad containing a lawn of oxygen nanosensors **(b)** The fluorescence emission intensity of oxygen nanosensors was compared under atmospheric and anoxic conditions using a confocal laser scanning microscope. The emission intensity of beads from six replicate slides was compared, with median atmospheric intensity of 766 intensity units (IQR = 425) and median anoxic intensity of 598 intensity units (IQR = 255). Statistical significance was calculated using a Mann-Whitney test (P < 0.01, ***) (N_External_ = 68, N_Internal_ = 87; acquired over 6 experimental replicates).

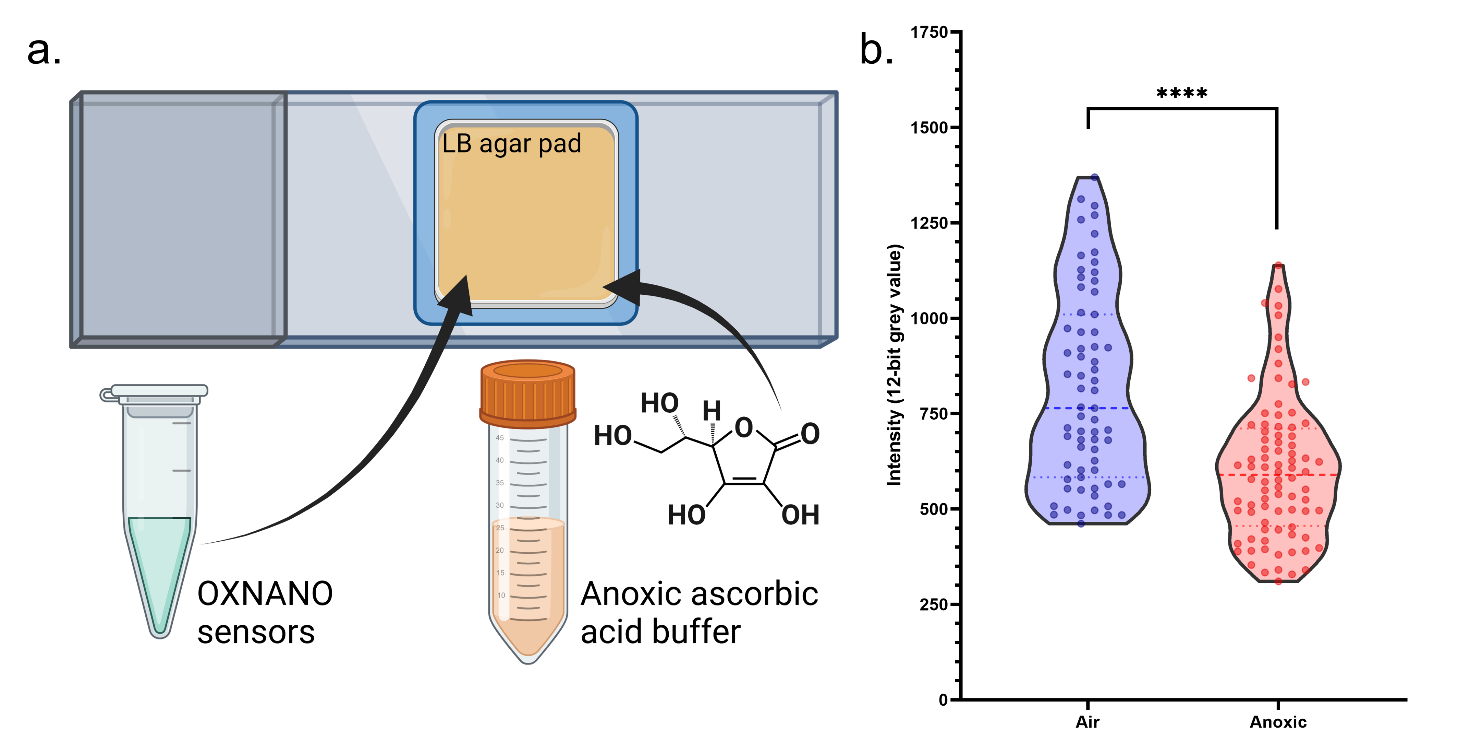

**Supplementary Figure 2. Methodology for colony biofilm thin sectioning.** A diagrammatic workflow for agarose stabilisation, fixation, paraffin embedding and thin sectioning of *E. coli* macrocolony biofilms. Specimens are embedded in cooled molten agarose, excised from a Petri dish and fixed in 4% (w/v) paraformaldehyde. Fixed blocks are then processed and paraffin embedded before microtome sectioning and mounting for optical imaging.

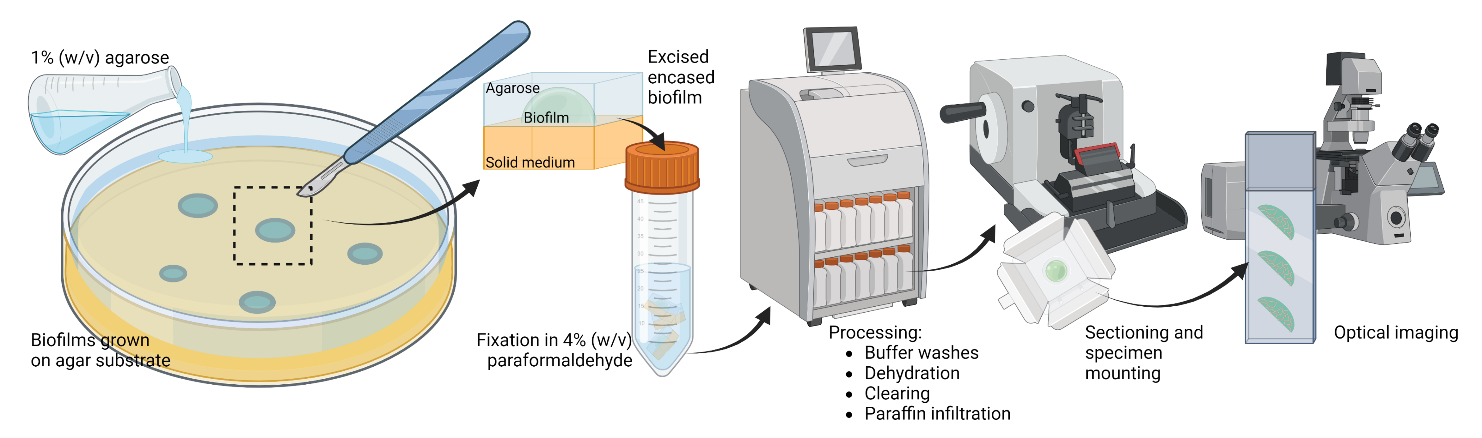

| Strain/Plasmid ID | Genotype | Source |
| --- | --- | --- |
| JM105 (DSMZ-3949) | Wild type  *endA1 glnV44 sbcB15 rpsL thi-1* Δ(*lac-proAB*) [F' *traD36 proAB*^+^ *lacI^q^ lacZ*Δ*M15*] *hsdR4*(*r_K_^-^m_K_* ^+^) | DSMZ, Germany |
| JM105 miniTn7::*HcRed* | Gm^R^ P._A1/04/03_::*HcRed* | ^66^ |
| JM105 miniTn7::*gfp* | Gm^R^ P._A1/04/03_::*gfp* | ^66^ |
| NEB-5alpha | *fhuA2*Δ(*argF-lacZ*)*U169 phoA glnV44* Φ*80*Δ(*lacZ*)*M15 gyrA96 recA1 relA1 endA1 thi-1 hsdR17* | New England Biolabs, USA |
| pAW9 | Oxygen reporter plasmid  Cm^R^ P*_cco2_*::*gfp* | ^28^ |

**Supplementary Table 1. List of bacterial strains and plasmids.**
